# The elusive actin cytoskeleton of a green alga expressing both conventional and divergent actins

**DOI:** 10.1101/554279

**Authors:** Evan W. Craig, David M. Mueller, Miroslava Schaffer, Benjamin D. Engel, Prachee Avasthi

**Affiliations:** Department of Anatomy and Cell Biology, University of Kansas Medical Center; Department of Molecular Structural Biology, Max Planck Institute for Biochemistry; Department of Ophthalmology, University of Kansas Medical Center

## Abstract

The green alga *Chlamydomonas reinhardtii* is a leading model system to study photosynthesis, cilia, and the generation of biological products. The cytoskeleton plays important roles in all of these cellular processes, but to date, the filamentous actin network within *Chlamydomonas* has remained elusive. By optimizing labeling conditions, we can now visualize distinct linear actin filaments at the posterior of the nucleus in both live and fixed vegetative cells. Using *in situ* cryo-electron tomography, we confirmed this localization by directly imaging actin filaments within the native cellular environment. The fluorescently-labeled structures are sensitive to the depolymerizing agent Latrunculin B (Lat B), demonstrating the specificity of our optimized labeling method. Interestingly, Lat B treatment resulted in the formation of a transient ring-like filamentous actin structure around the nucleus. The assembly of this perinuclear ring is dependent upon a second actin isoform, NAP1, which is strongly upregulated upon Lat B treatment and is insensitive to Lat B-induced depolymerization. Our study combines orthogonal strategies to provide the first detailed visual characterization of filamentous actins in *Chlamydomonas*, allowing insights into the coordinated functions of two actin isoforms expressed within the same cell.

## Introduction

Actin is a highly conserved protein that is found across eukaryotes and is essential for survival in most eukaryotic cells (Pollard et al., 2000). In yeast, the sole actin gene forms distinct structures such as actin patches, cables, and cytokinetic rings (Kilmarten and Adams 1984; Warren et al., 2002). These structures function in endocytosis, organelle and protein transport, and cell division, respectively. Unlike yeast, mammalian cells have six different actin isoforms encoded by six separate genes, some of which require complex interactions between actin isoforms, and collectively are involved in functions such as regulating cell morphology, motility, mechanotransduction, membrane regulation, and intracellular trafficking. Single actin knockout studies in mice revealed that loss of one gene causes a compensatory upregulation of a subset of the remaining actin isoforms (Perrin and Ervasti 2010). To understand the overlapping functions and co-regulation of multiple actin isoforms, we chose a model organism that expresses only two actin genes.

The genome of the unicellular green alga *Chlamydomonas reinhardtii* contains two actin genes that vary significantly in sequence. Inner dynein arm 5 (IDA5) is a highly conserved conventional actin, whereas novel actin-like protein 1 (NAP1) is a divergent actin that only shares ~65% sequence identity with mammalian actin (Kato-Minoura et al., 1997; Onishi et al., 2016). Functionally, genetic loss of *IDA5*, a condition in which NAP1 is expressed at low levels, results in slow swimming (Ohara et al., 1998) and early ciliary growth defects due to reduced ciliary protein trafficking (Avasthi et al., 2014). Ciliary growth ultimately proceeds and can reach normal length. However, when both actins are disrupted acutely, *Chlamydomonas* cells show dramatic defects in ciliary protein synthesis, vesicular trafficking, and organization of a key gating region dictating ciliary protein composition (Jack et al., 2018). Given that *ida5* mutants expressing NAP1 alone don’t show these defects, it appears NAP1 can largely perform actin-dependent functions needed for ciliary assembly despite its sequence divergence with IDA5. Although we have been able to genetically and chemically dissect the functions of the individual actin isoforms, detailed visual characterization of filamentous actin networks has eluded the field.

While actin filaments are readily visualized by traditional phallotoxin staining in mammalian systems, a variety of protein and cellular differences complicate actin visualization in protists. In the parasite *Toxoplasma gondii*, the actin gene *ACT1* shares 83% sequence identity with mammalian actin and is required for cell motility, yet filamentous actin is undetectable by phalloidin staining (Dobrowolski et al., 1997). Other conventional filamentous actin labeling techniques such as Life-Act and SiR-actin fail to label filamentous actin in this parasite (Periz et al., 2017), demonstrating that that actin labeling optimization is needed for unique systems. This may be due to highly dynamic filament turnover and unique actin polymerization kinetics (Miraldi et al., 2008) exemplified in *Toxoplasma gondii* and closely related *Plasmodium falciparum.* Biochemical assays also show 97% of the parasites’ actin exists in globular form (Dobrowolski et al., 1997; Skillman et al., 2011; Wetzel et al., 2003), creating unfavorable conditions for filamentous actin staining. As in parasites, *Chlamydomonas* actin visualization with conventional strategies has also been challenging. Actin antibodies do not discriminate between filamentous and monomeric actin, and previous attempts to visualize the filamentous actin cytoskeleton using fluorescent phallotoxins resulted in diffuse signal throughout the cytoplasm in vegetative *Chlamydomonas* cells (Harper et al.,1992), leading some to conclude that these cells made few, if any, filaments (Harper et al.,1992). The only condition where phallotoxins have been shown to clearly label filamentous actin in *Chlamydomonas* is in gametes, where filamentous actin-rich tubules can be seen at the apical surface between flagella upon mating or artificial induction (Detmers et al., 1985).

Advancements in vegetative *Chlamydomonas* actin filament visualization came from live cell imaging using strains expressing the fluorescently-tagged filament binding peptide, LifeAct (Avasthi et al., 2014; Onishi et al., 2016). This method identified linear structures at the posterior of the nucleus and less frequently at the base of the flagella. Acute treatment with the actin depolymerizing agent Latrunculin B (Lat B) eliminated this labeling (Avasthi et al. 2014; Onishi et al., 2016), demonstrating the specificity of the fluorescent LifeAct for actin filaments. However, longer incubation with Lat B restored partial signal with a slightly different distribution (Onishi et al., 2016). This new signal likely represents the Lat B-induced upregulation of a second *Chlamydomonas* actin normally expressed at low levels, the novel actin-like protein NAP1 (Onishi et al., 2016; Kato-Minoura et al., 1998). Despite the identification of a clear filamentous actin-based perinuclear structure using fluorescent LifeAct, technical challenges to maintaining LifeAct expression and the inability to preserve labeled structures for co-localization with cellular organelles in fixed cells required a different method of actin visualization.

We reasoned that *Chlamydomonas* actin, which shares 90% sequence identity with mammalian actins, is inherently capable of binding fluorescent phallotoxins due to the intense staining of fertilization tubules in gametes. In this study, we developed an optimized protocol for phalloidin staining that recapitulated LifeAct labeling. Using this method, and corroborating with live cell visualization and cryo-electron tomography (cryo-ET), we can now show for the first time how actin filaments are localized and redistributed in vegetative and gametic *Chlamydomonas* cells. In addition, we applied this staining method to mutants of each actin isotype to reveal new insights into isoform-specific organization and function.

## Methods

### Phalloidin staining

Cells were grown in 2 mL of TAP (tris acetate phosphate) liquid media on a roller drum for 17 hr (overnight). It is essential to select healthy cells by centrifuging 1 mL of cell culture at 1800 RPM for 1.5 min. The supernatant was discarded and cells were resuspended in 600 μL fresh TAP. 200 uL of resuspended cells were adhered on Poly-L-Lysine-coated coverslips for 5 min and covered. The liquid was tilted off the coverslips and replaced with 4% fresh paraformaldehyde (PFA) in 10 mM HEPES pH 7.24. The pattern of staining changes (producing bright pyrenoid signal) and non-specific cytoplasmic fluorescence increases if using PFA that is not fresh. Coverslips were washed with 1x PBS for 3 min. For cell permeabilization, coverslips were submerged in a coplin jar containing 80% pre-cooled acetone and then incubated for 5 min at – 20° C. Coverslips were quickly transferred to a second coplin jar containing 100% pre-cooled acetone and incubated again for 5 min at −20° C. Coverslips were allowed to air dry for a minimum of 2 min, and for longer if needed. Next, cells were rehydrated by transferring coverslips to a coplin jar containing 1x PBS for 5 min. Coverslips were then stained with Atto 488 Phalloidin (49409, Sigma) for 16 min in the dark, significantly reducing background and increasing the signal to noise ratio. The Atto 488 Phalloidin reagent greatly enhanced photostability and brightness compared to Alexa-488, which allowed for more uniform and reproducible filament labeling. Phalloidin incubation was followed by a wash in 1x PBS for 5 min. Finally, coverslips were mounted on slides with self-sealing Fluoromount-G™ (Invitrogen) as quickly as possible.

### Cell vitrification and FIB milling

*Chlamydomonas* cells were grown and prepared for cryo-ET as previously described (Bykov et al., 2017; Albert et al., 2017). We used the *mat3-4* strain (Umen and Goodenough, 2001), which has smaller cells that improve vitrification by plunge-freezing. Cells were grown with constant light (~90 μmol photons m^-2^ s^-1^) in Tris-acetate-phosphate (TAP) medium, bubbling with normal air. Liquid culture (5 μL, ~1000 cells/μL) was blotted onto carbon-coated 200-mesh copper grids (Quantifoil Micro Tools), which were then plunged into a liquid ethane-propane mixture using a Vitrobot Mark 4 (FEI Thermo Fisher). Frozen grids were mounted into modified Autogrids (FEI Thermo Fisher) and loaded into either a Scios (FEI Thermo Fisher) or a Quanta 3D FEG (FEI Thermo Fisher) FIB/SEM microscope. The grids were coated with thin layer of organometallic platinum using the gas injection system (GIS, FEI Thermo Fisher), and then the Ga+ beam was used to mill thin lamellae (100-200 nm thick), following the protocols detailed in (Schaffer et al., 2015; Schaffer et al., 2017).

### Cryo-electron tomography

Tomography was performed on a 300 kV Titan Krios (FEI Thermo Fisher) equipped with a Quantum postcolumn energy filter (Gatan) and a K2 Summit direct electron detector (Gatan) operated in movie mode at 12 frames per second. Using SerialEM software (Mastronarde, 2005), tilt series were acquired over an angular range of approximately −60° to +60°, using a bidirectional tilt scheme with 2° increments. Cumulative electron dose was ~100 e-/Å^2^, the object pixel size was 3.42 Å, and defocus ranged from −4 to −6 μm. Raw image frames were aligned with MotionCorr2 (Zheng et al., 2017) software and tomograms were reconstructed in IMOD software (Kremer et al, 1996) using patch tracking and back projection. The images displayed in this paper were binned twice. To enhance the contrast, we used the tom deconv deconvolution filter (https://github.com/dtegunov/tom_deconv).

### Fertilization tubule induction

To induce gametogenesis, cells were grown in M-N (M1 minimal media without nitrogen) overnight under growth lighting. Gametes were mixed with dibuteryl cAMP (13.5mM) and papaverine (135μM) for 45 minutes to induce fertilization tubule formation (modified from Wilson et al., 1997). Cells were then stained in accordance with our phalloidin protocol (see above).

### Latrunculin B treatment

Latrunculin B was purchased from Sigma and used with the specified concentrations and incubation times. Typically, depolymerizing treatment was performed for 10 minutes, and cells were treated for 45 minutes to induce NAP1 upregulation and ring formation.

### Chlamydomonas strains

The wild-type strain CC-125 mt + and *mat3-4* strain CC-3994 mt + were obtained from the Chlamydomonas Resource Center (University of Minnesota). The *nap1* mutant was a generous gift from Fred Cross (The Rockefeller University), Masayuki Onishi (Stanford University), and John Pringle (Stanford University). The LifeAct-Venus wild-type transformant was gifted by Masayuki Onishi.

## Results

### Filamentous actin visualization in vegetative Chlamydomonas achieved by an optimized phalloidin staining protocol

To optimize phalloidin labeling, which previously produced only weak, diffuse, seemingly nonspecific signal in vegetative cells (Harper et al., 1992) (**Figure 1A, C**), we used a combination of bright and photostable Atto fluorophores and reduced incubation times to limit background fluorescence. We also improved the imaging by employing deconvolution microscopy to remove out of focus light. Using optimized protocols (see methods), we found that the pattern of phalloidin staining matched what was seen for LifeAct-Venus fluorescence, with a tangle of linear filaments at the posterior of the nucleus (**Figure 1B, D, F**), providing further support that the previously identified mid-cell LifeAct signal (Avasthi, et al 2014; Onishi, et al. 2016) represents the filamentous actin population. Phalloidin-labeled structures were disrupted when treated with the actin-depolymerizing drug, Latrunculin B (Lat B) (**Figure 2C**).

**Figure 1.**
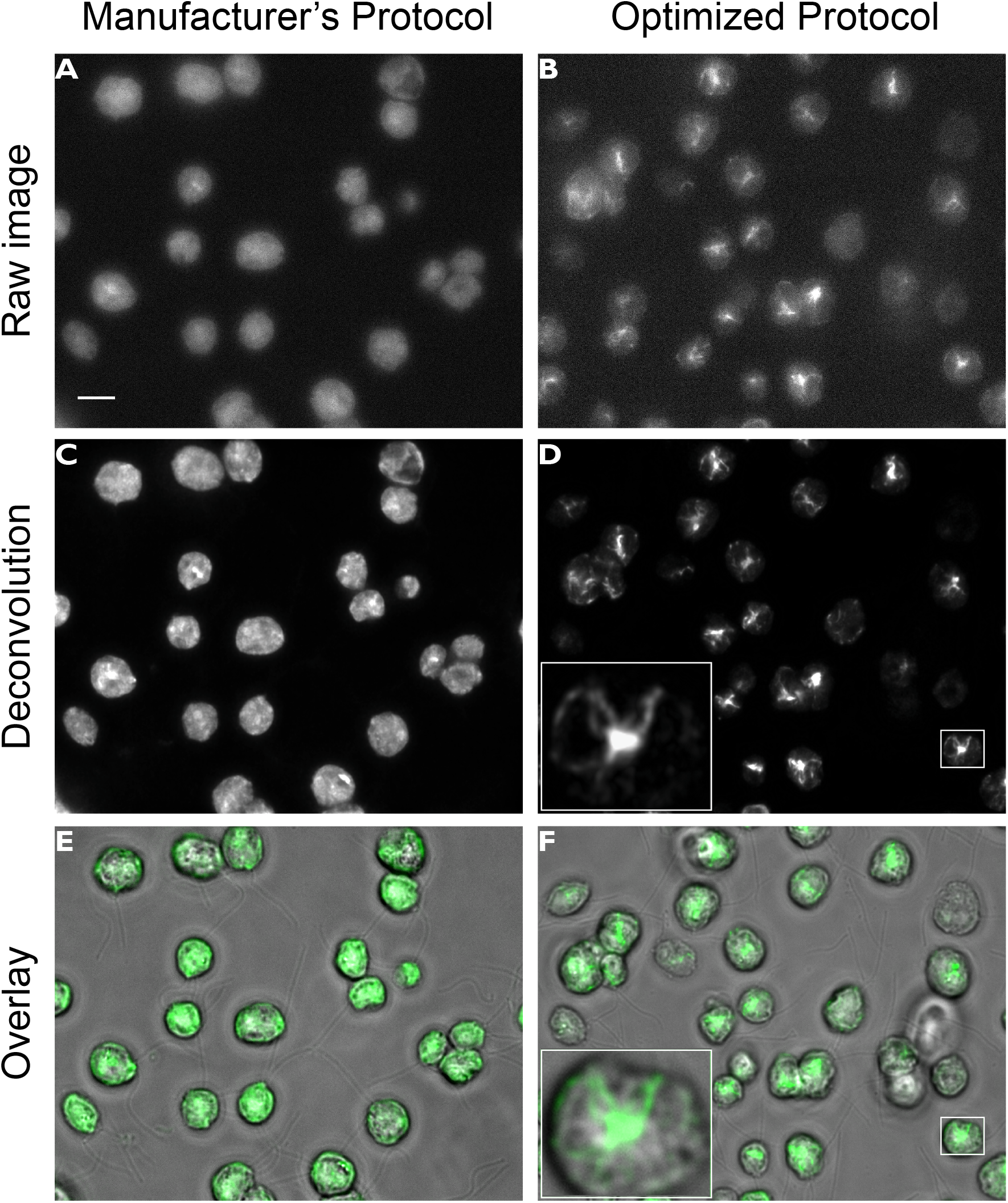
Filamentous actin staining optimization using phalloidin in *Chlamydomonas reinhardtii.* **A)** Raw fluorescence image of phalloidin-stained vegetative *Chlamydomonas* cells using the manufacturer’s recommended protocol and Alexa Fluor 488 Phalloidin. Signal is generally bright with hazy fluorescence throughout the cell, similar to previous reports. **B)** Raw fluorescence image using our optimized phalloidin protocol and Atto 488 Phalloidin (49409, Sigma) reagent. Signal from filamentous actin is clearly present. **C)** Deconvolution of the image in A does not reveal much actin signal that can be easily distinguished from the high background fluorescence. **D)** Deconvolution of B shows filamentous actin posterior of the nucleus and filaments spanning across the cell body. **E&F)** Overlay of brightfield and fluorescence channels with phalloidin signal in green. In vegetative cells, the brightness and staining consistency were greatly enhanced by using the Atto 488 conjugate instead of Alexa Fluor 488. Scale bar is 5 μm.

**Figure 2.**
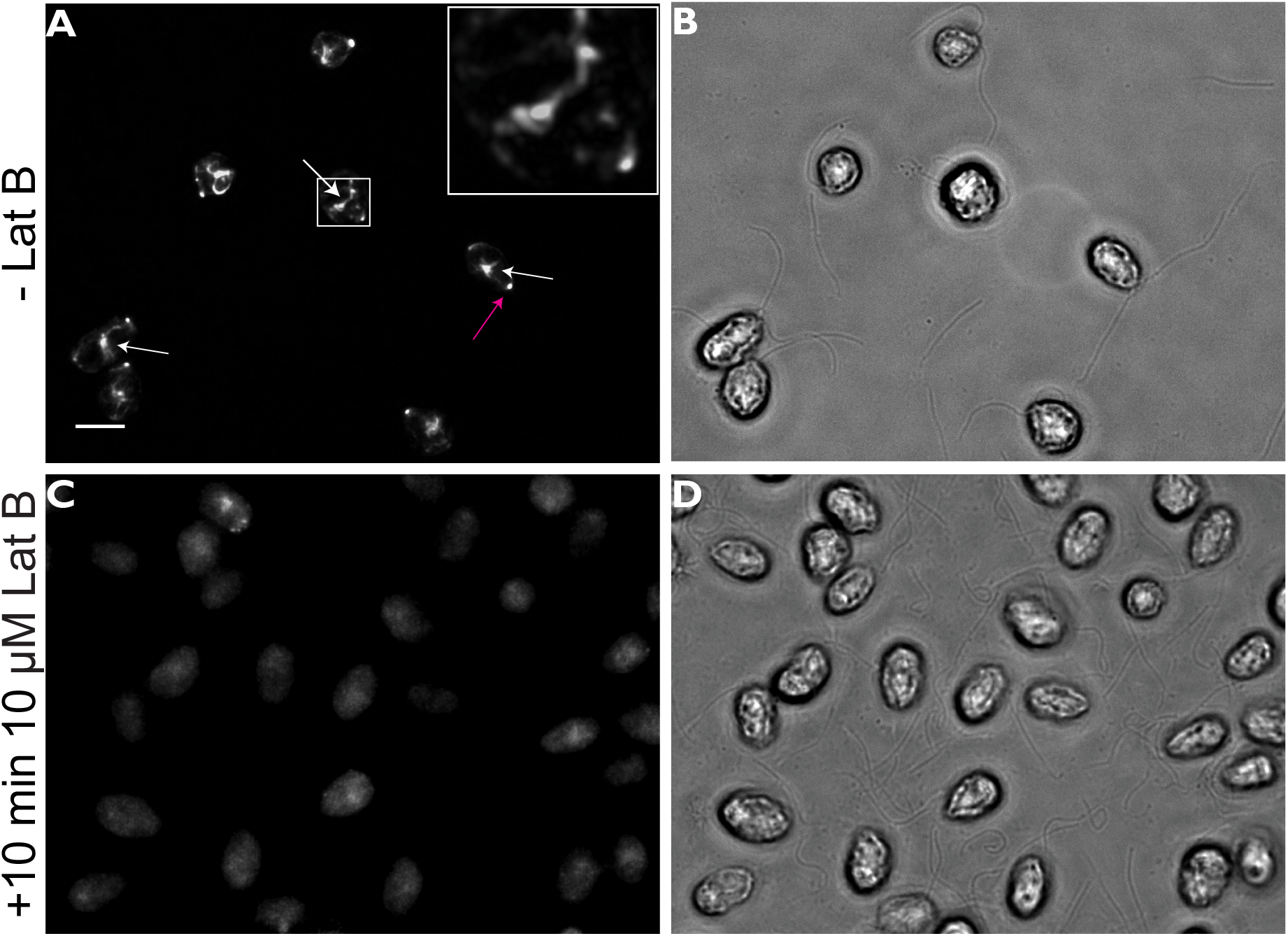
Phalloidin-labeled filamentous actin depolymerizes upon Latrunculin B treatment in wild-type CC-125 cells. **A)** Gametic CC-125 cells stained with Atto 488 Phalloidin, showing mid-cell actin staining (white arrows) and apical actin fluorescence (magenta arrow). **C)** Atto 488 Phalloidin-stained gametic CC-125 cells after 10 minutes of treatment with 10 μM Latrunculin B. Filamentous actin signal dramatically decreases. **B&D)** Brightfield images show filamentous actin signal in relation to the cell body and flagella. Scale bar is 5 μm.

### In situ cryo-electron tomography of actin filaments

To directly visualize actin filaments within the native cellular environment, we rapidly froze vegetative *Chlamydomonas* cells in vitreous ice, thinned these cells by focused ion beam milling, and then imaged them by cryo-ET (Asano et al., 2016). Helical filaments with a diameter of ~7 nm (consistent with filamentous actin) were observed in the cytosol at the posterior side of the nucleus, near the nuclear envelope, ER, and Golgi (**Figure 3**). In addition to relatively straight individual actin filaments (**Figure 3A-B**), we also observed loosely tangled bundles of filaments with increased local curvature (**Figure 3C**).

**Figure 3.**
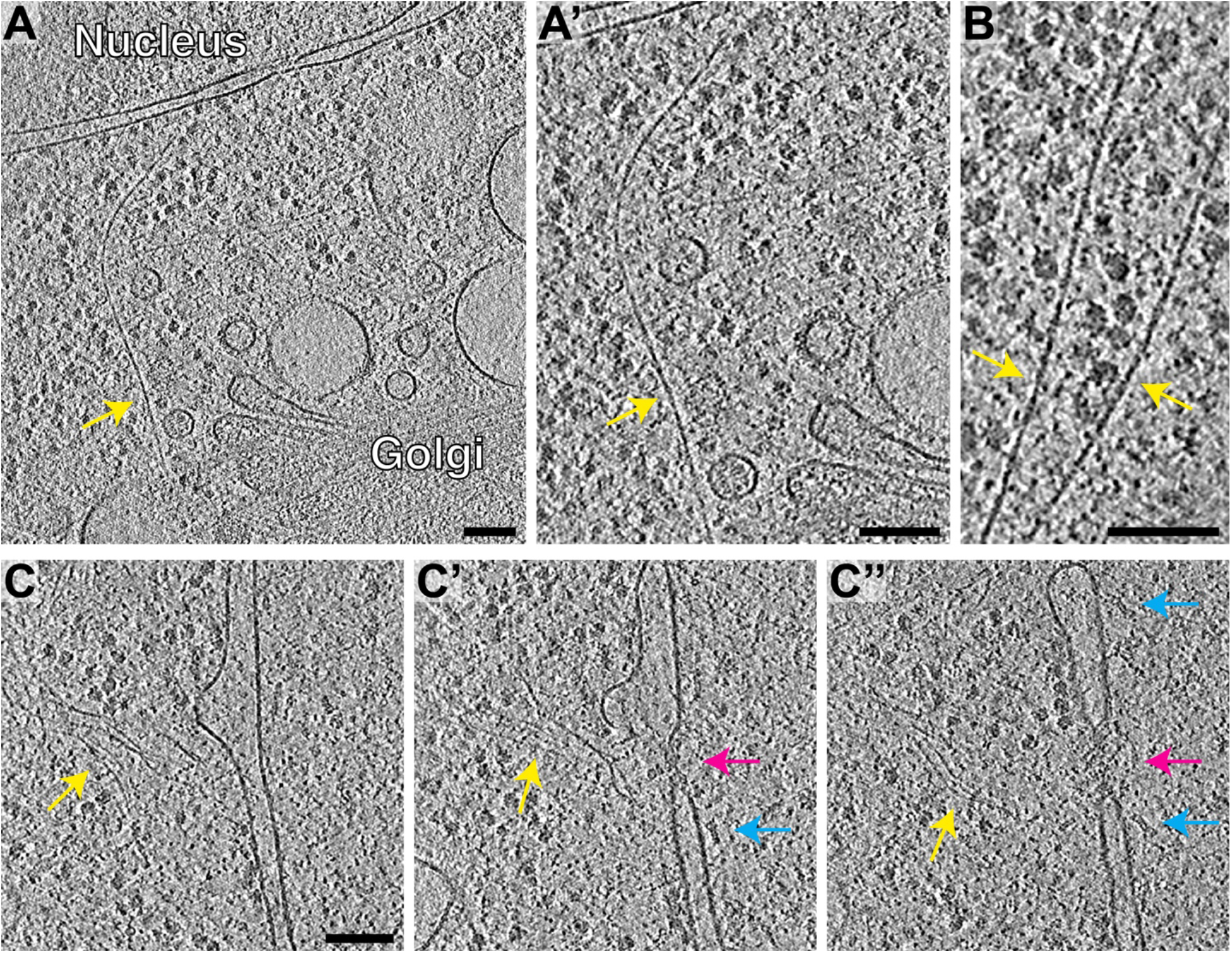
Direct visualization of actin filaments in *Chlamydomonas reinhardtii* by *in situ* cryo-ET. **A-B)** Slices through tomographic volumes. Yellow arrows indicate individual actin filaments near the nuclear envelope and Golgi apparatus. **A**’ is an enlarged view of the filament from **A**. The helical structure of the actin filaments is apparent in **B. C)** A loose bundle of actin filaments (yellow arrow) in close proximity to a nuclear pore complex (magenta arrow). Clearly resolved nuclear proteasomes tethered to the nuclear pore complex (blue arrows, previously described in Albert et al., 2017) illustrate the molecular clarity of this tomogram. **C, C**’, and **C**” show three different slices through one tomographic volume. Scale bars are 100 nm in all panels.

### Actin dynamics in vegetative and gametic cells

Filamentous actin localization appears to differ between cell states in *Chlamydomonas reinhardtii.* When stained with Atto 488 Phalloidin, mid-cell filamentous signal was observed near the posterior of the nucleus in both vegetative and gametic cells. However, compared to vegetative cells (**Figure 4A-C**), the perinuclear signal in non-cycling gametes became prominent and uniform throughout the cellular population (**Figure 4D-F**), likely due to the strong upregulation of actin expression (Ning et al., 2013). Interestingly, gametic cells often showed a single apical fluorescent spot between the two flagella (**Figure 4F, inset**). In the case of vegetative cells, this apical fluorescence was observed much less frequently and formed one, two, three, or four spots; this variability may be attributed to working with an asynchronous cell population.

**Figure 4.**
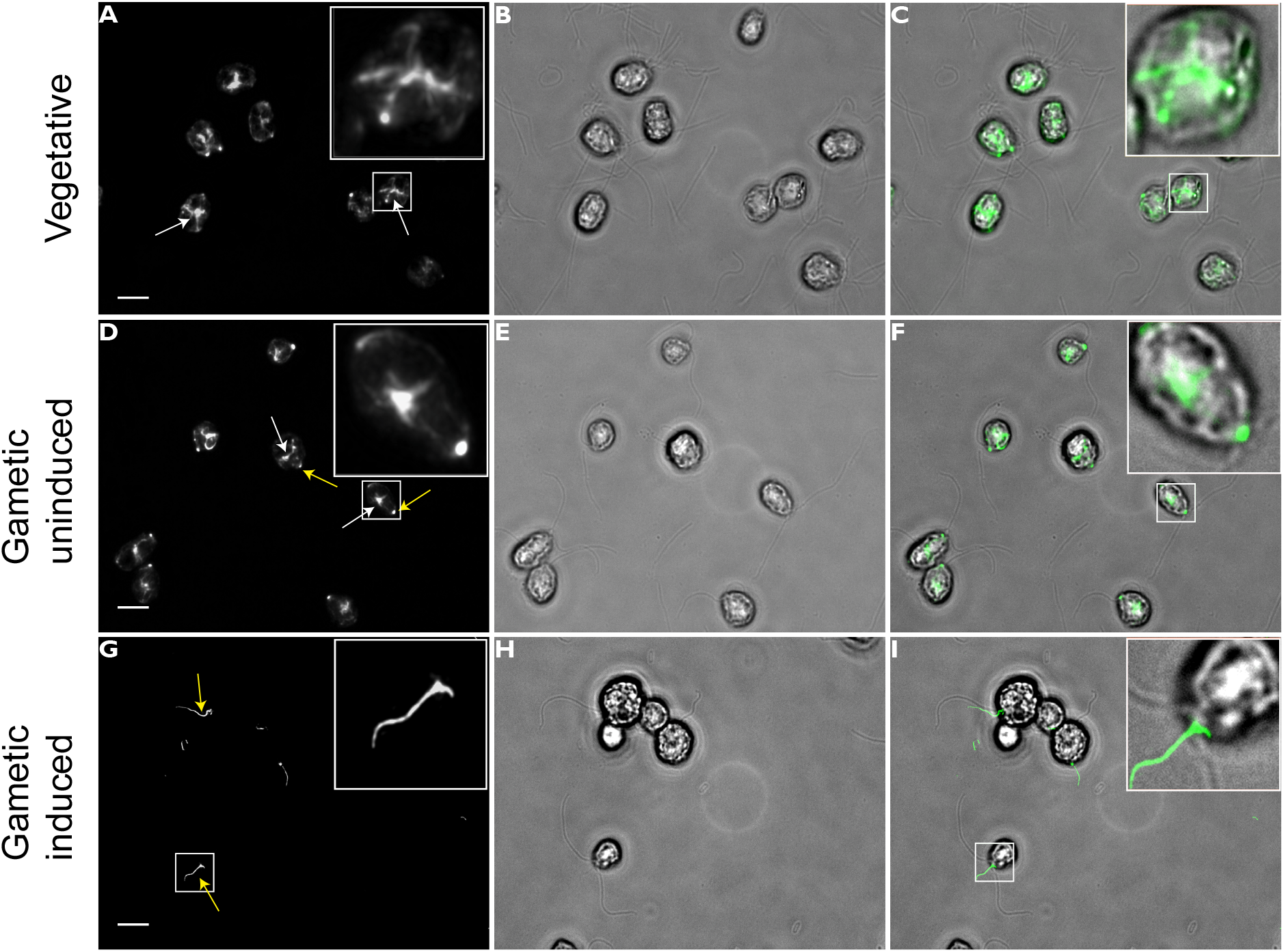
Filamentous actin localization in *Chlamydomonas* cell types. **A)** Filamentous actin (white arrows) localizes posterior of the nucleus in vegetative wild-type CC-125 cells. **D)** Filamentous actin in CC-125 gametes show mid-cell localization similar to vegetative cells, but also contain an actin population localizing between the flagella (yellow arrows). **G)** Induced gametes stained with Atto 488 Phalloidin display actin-rich fertilization tubule structures. **B, E, & H** are single slice brightfield images. **C, F, & I** are overlays of brightfield and fluorescence channels, with filamentous actin signal in green. Scale bar is 5 μm.

The apical filamentous actin foci may mark the location of eventual fertilization tubule formation prior to induction. During artificial induction of gametes, fertilization tubules form after the addition of papavarine and dibutyryl cAMP. This actin-rich fusion organelle protrudes from the apex of the cell, the same location where we observed apical actin accumulation in uninduced gametes (**Figure 4I, inset**). During tubule induction, the mid-cell filamentous actin population disappeared (compare **Figures 4F and 4I**). Based upon this loss of mid-cell actin and previous transcriptomic studies showing a 2.4-fold increased expression of the actin depolymerizing factor cofilin when fertilization tubules are induced (Ning et al, 2013), we hypothesize that the perinuclear actin filaments are depolymerized and repurposed to build the filamentous actin-rich fertilization tubule.

### Visualization of conventional and non-conventional actins

We examined the cellular distribution of the two *Chlamydomonas* actins independently by staining *nap1* mutants (expressing only IDA5) and *ida5* mutants (expressing only NAP1) with Atto 488 Phalloidin. Atto 488 Phalloidin staining in *nap1* vegetative cells displayed wild-type-like mid-cell actin signal (**Figure 5A, 5B**), and a single apical spot was present in almost every cell (magenta arrows, **Figure 5B**). However, in *ida5* mutants, signal remained weak and hazy (**Figure 5C**). This may be due to low NAP1 abundance or slight sequence differences in the phalloidin binding sites of IDA5 and NAP1. Actin has two methionine residues that are critical for phalloidin binding, and these are conserved in the IDA5 sequence. However, NAP1 contains mutations at both methionine sites, which could alter phalloidin’s ability to label NAP1 filaments (Vandekerckhovel et al., 1985).

**Figure 5.**
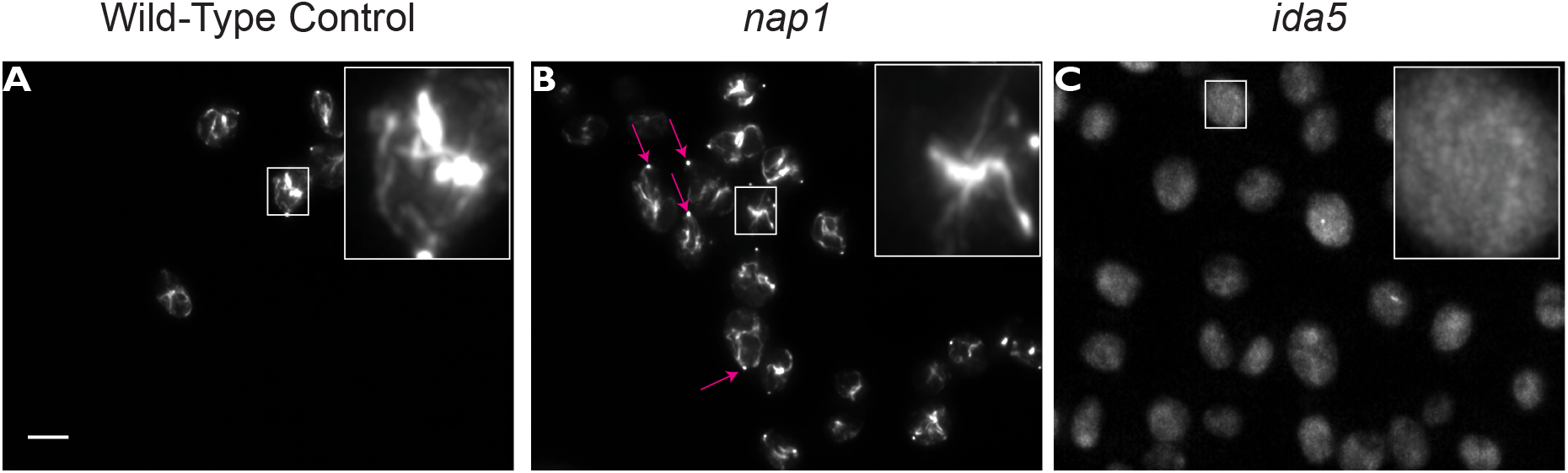
Phalloidin staining of *ida5* and *nap1* mutants. **A)** Vegetative wild-type CC-125 cells stained with Atto 488 Phalloidin. **B)** Vegetative *nap1* cells (expressing only IDA5) stained with Atto 488 Phalloidin display wild-type-like mid-cell actin signal and frequent apical signal (magenta arrows). **C)** Vegetative *ida5* cells (expressing only NAP1) stained with Atto 488 Phalloidin shows non-specific signal. Scale bar is 5 μm.

Treating wild-type cells with the actin-depolymerizing drug Lat B for 10 minutes eliminated the specific mid-cell filamentous actin staining (**Figure 6B**). Lat B incubation, which is known to upregulate NAP1 expression (Onishi, 2018), resulted in the formation of a transient ring structure around nuclei (**Figure 6C**), appearing at ~45 minutes of Lat B treatment and dissipating by two hours. Live cell imaging using a LifeAct-Venus transformant revealed a similar ring phenotype, but rings seemed to occur in fewer cells using this filamentous actin probe. (**Figure 6D**). Based on the Lat B-induced upregulation of NAP1 expression and the insensitivity of NAP1 to Lat B (Onishi et al., 2016), we hypothesized that this ring structure is comprised of NAP1 filaments. To test this, we performed a 45 minute Lat B treatment on the *nap1* mutant background. Atto 488 Phalloidin-stained *nap1* cells displayed similar wild-type-like mid-cell actin signal. (**Figure 7E**). However, Lat B treatment did not result in the formation of filamentous ring structures in *nap1* cells (**Figure 7G**), suggesting this actin population is NAP1-dependent.

**Figure 6.**
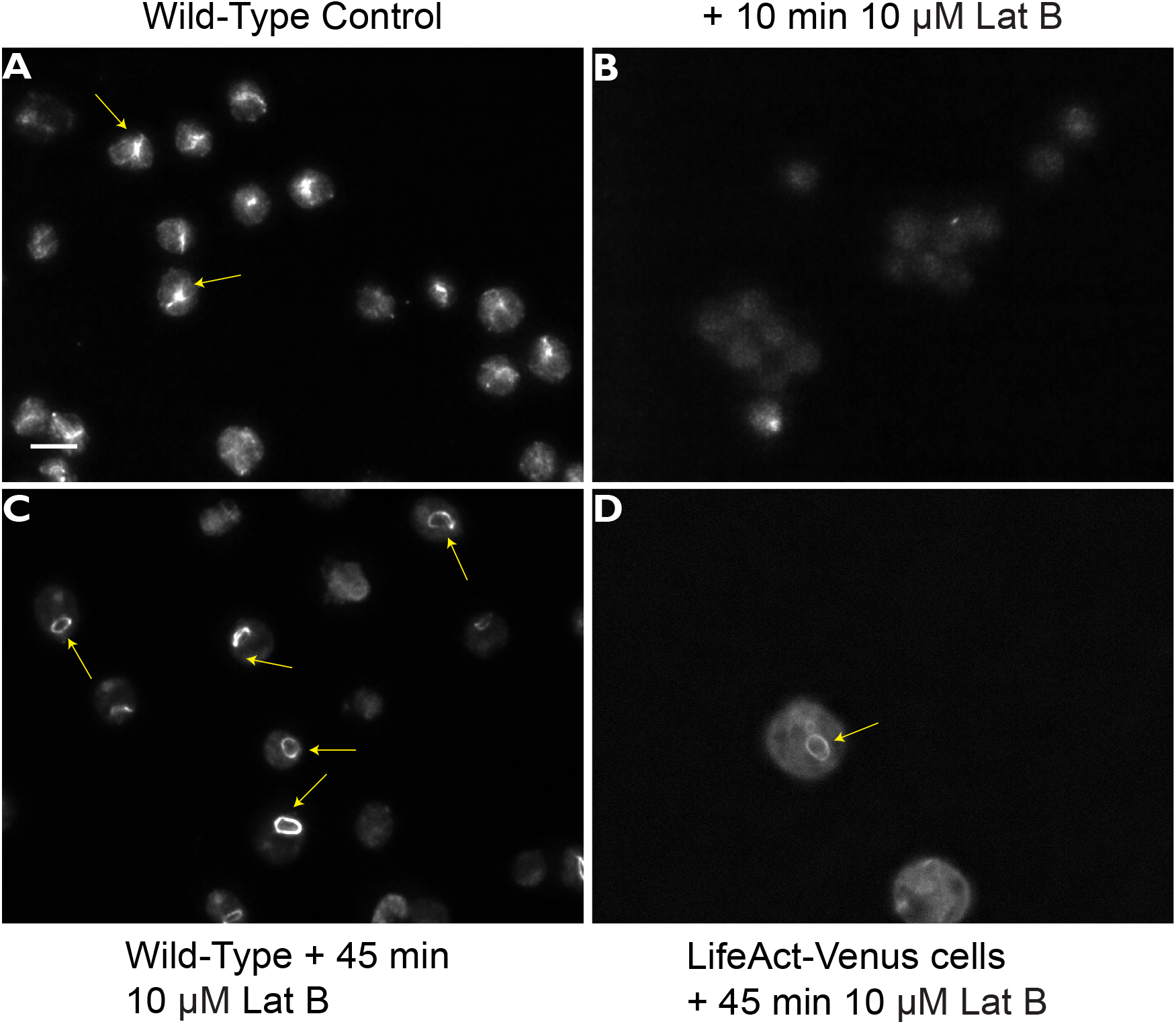
Ring-like actin structures in CC-125 cells are induced by prolonged Latrunculin B treatment. **A)** Vegetative wild-type CC-125 cells stained with Atto 488 Phalloidin. Yellow arrows show the filamentous actin signal. **B)** The filamentous actin signal disappears after 10 minutes of treatment with 10μM Lat B. Cells stained with Atto 488 Phalloidin. **C)** Yellow arrows show filamentous actin ring structures in many CC-125 cells after 45 minutes of treatment with 10 μM Lat B. Cells stained with Atto 488 Phalloidin. **D)** Live LifeAct-Venus cells treated with 10 μM LatB for 45 minutes also show the actin ring phenotype. Scale bar is 5 μm.

**Figure 7.**
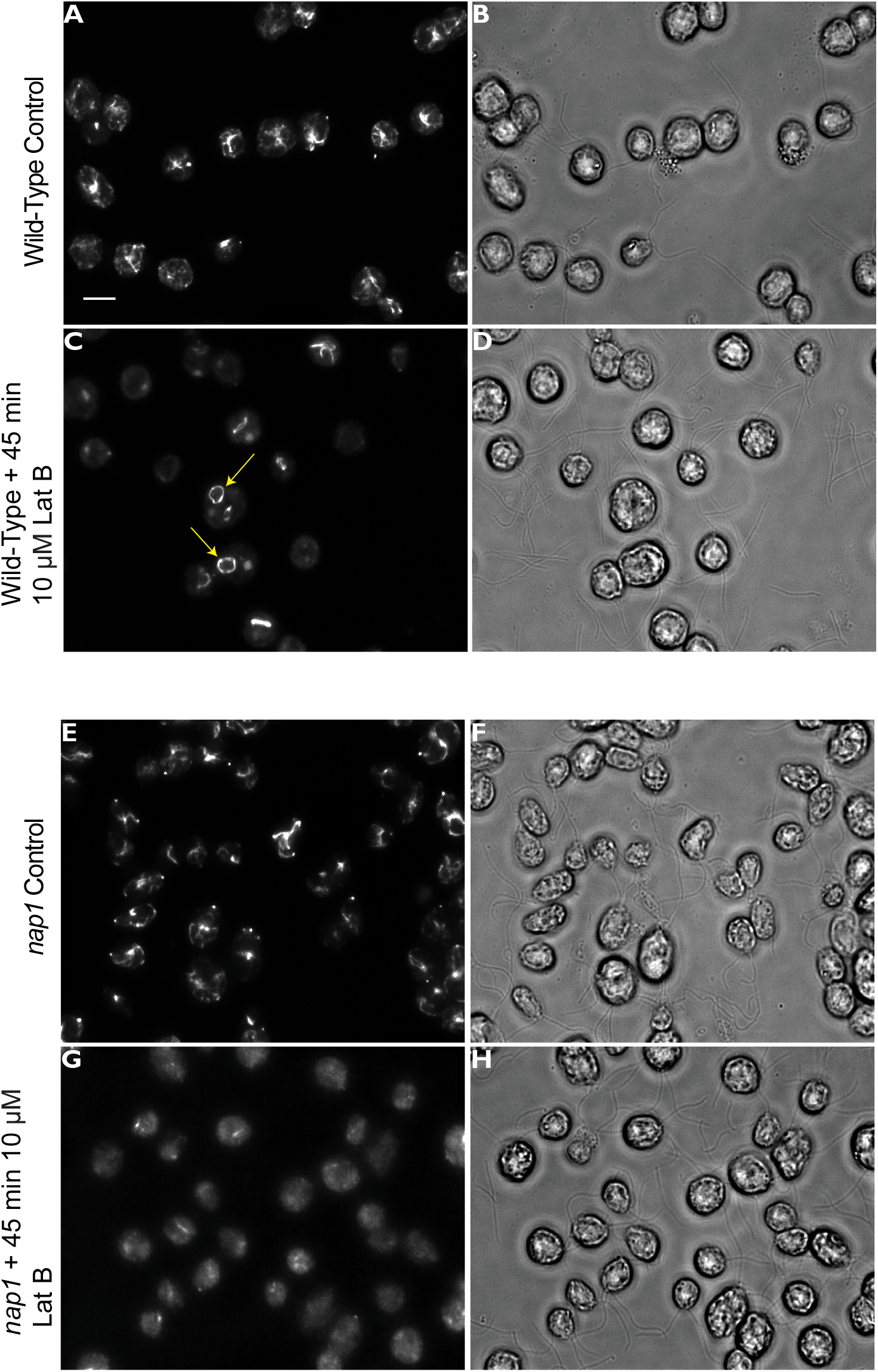
The ring-like filamentous structure in Lat B-treated cells is likely composed of NAP1 filaments. **A)** Vegetative wild-type CC-125 cells stained with Atto 488 Phalloidin, showing the typical midcell actin signal. **C)** Atto 488 Phalloidin-stained vegetative CC-125 cells treated with Lat B for 1 hour. Filamentous actin is deploymerized. **E)** Atto 488 Phalloidin-stained vegetative *nap1* mutant cells show similar mid-cell staining compared to wild type. **G)** Atto 488 Phalloidin-stained vegetative *nap1* mutant cells treated with Lat B for 1 hour do not show an filamentous actin ring signal. Scale bar is 5 μm.

## Discussion

The coordinated expression and polymerization of two actins that share ~65% sequence identity within the same cell can provide broader insight into mechanisms of actin regulation and segregation of actin-dependent functions *in vivo.* The two *Chlamydomonas* actin genes are poorly characterized. One reason for this is that the *ida5* null mutant appears to have a mild, wild-type-like phenotype. We now believe this is due to compensation by the upregulation of NAP1 in *ida5* mutants (Kato Minoura, 2005; Onishi et al., 2016; Onishi et al., 2018). Loss of both actins is lethal (Onishi et al., 2016), and our recent studies demonstrate critical functions for at least one actin at multiple stages of ciliary assembly (Jack et al., 2018).

Outside of the compensatory functions, the requirement for dual expression is not clear since NAP1 expression is undetectable under normal conditions and only upregulated slightly in the absence of IDA5 or strongly upon flagellar reassembly following deflagellation (Hirono et al., 2003). Despite the upregulation of NAP1 upon flagellar regeneration, we previously found that *nap1* null mutants have no flagellar assembly defect outside of the inability to buffer perturbations to IDA5 (Jack et al., 2018). However, NAP1 can largely perform the functions of filamentous IDA5, as *ida5* mutants show only slow swimming. This is likely due to the loss of four axonemal inner dynein arm subspecies within flagella (Kato-Minoura et al., 1997) and slow initial phases of flagellar assembly (Avasthi et al., 2014). Mating type plus gametes of the *ida5* mutant cannot form structurally normal actin-rich fertilization tubules, resulting in a drastically reduced mating efficiency (Kato-Minoura et al., 1997).

To complement our functional studies of IDA5 and NAP1, we wanted to understand if the two proteins exhibit similar localization and dynamics. However, actins in *Chlamydomonas* have been difficult to visualize by fluorescence microscopy and traditional electron microscopy. Actin mutant phenotypes suggest actin localization in flagella, fertilization tubules, and the cleavage furrow. Indeed, this has been visualized using phalloidin in fertilization tubules (Wilson et al., 1997) and actin antibodies (which cannot discriminate between actin monomers and filaments) in the cleavage furrow (Ehler and Dutcher 1998; Kato-Minoura et al., 1998). It was reported that phalloidin did not clearly label filamentous structures in vegetative *Chlamydomonas* cells after two hours of staining; instead, only bright fluorescence throughout the cell body could be observed (Harper et al., 1992). We initially encountered similar results, even when using a much shorter incubation time of 30 minutes. However, reducing the staining time to 16 minutes resulted in optimal signal to noise and clear labeling of filamentous structures. Despite this improvement, many phalloidin trials using this 16-minute incubation time resulted in bright and diffuse signal in the cell body, and cells were susceptible to rapid photobleaching during imaging. These inconsistencies were alleviated after switching to a different phalloidin conjugate, Atto 488 Phalloidin, which produced a clear and consistent photostable signal. Electing for this reagent, combined with our optimized actin visualization protocol, lead to a powerful and reproducible method for visualizing filamentous actin in fixed vegetative *Chlamydomonas* cells. Combining three visualization methods (live cell LifeAct labeling, fixed cell phalloidin labeling, and direct imaging of filaments by cryo-ET) with analysis of Lat B sensitivity, our study confirms that *Chlamydomonas* cells have a *bona fide* actin filament network at the posterior of the nucleus and occasionally at the apical cell surface.

Our study reveals previously unreported filamentous structures in *Chlamydomonas.* Most obvious is the array of filamentous actin localized posterior of the nucleus in vegetative and gametic cell types. The actin-binding and nucleating proteins responsible for coordinating this actin population are not known, but recent findings in our lab strongly suggest a role for actin in trafficking vesicles from the Golgi (Jack et al., 2018). This mid-cell actin revealed by Atto 488 Phalloidin labeling, along with Cryo-ET showing distinct actin filaments near the Golgi (**Figure 3A**), steers us to investigate the functional role of actin in ciliary protein transport, which we believe is myosin-dependent (Avasthi et al., 2014). The mid-cell actin is likely composed of IDA5 filaments, as we see similar actin localization when labeling *nap1* mutant cells and no filamentous signal in this region in when labeling *ida5* mutants. NAP1 expression is significantly lower in unperturbed vegetative cells, but increased in *ida5* mutant cells (Ohara et al., 1998). While we have struggled to visualize this population of filamentous NAP1 actin with phalloidin staining, we are pursuing improved stability LifeAct probes and alternative strategies.

We consistently observed apical accumulation of filamentous actin between the flagella in gametic cells, and more variable apical signal appearance in vegetative cells. We believe this filamentous actin population in gametes serves as an actin recruitment site, perhaps supplying a precursor pool of actin monomers for building the highly filamentous fertilization tubule formed in mating type plus gametic cells. It was previously unknown whether actin was locally assembled at the site of fertilization tubule formation, perhaps from newly synthesized actin as actin expression is known to increase during gamete activation (Ning et al, 2013). However, we note the absence of the mid-cell actin when we label induced gametes, suggesting fertilization tubule actin may instead be remodeled from the mid-cell population.

We rarely observed actin mislocalization from the sub-flagellar region or multiple spots of apical accumulation in gametes. A well-conserved function of actin is in endocytosis. It is possible that the variable spots we see at the cell periphery in vegetative cells are analogous to endocytic patches/pits. While there is little evidence for endocytosis in *Chlamydomonas*, studies of cell adhesion during mating show that activity-dependent redistribution of signaling proteins from the cell body plasma membrane into cilia seems to occur in a microtubule-dependent manner but is independent of ciliary motors (Belzile et al., 2013). This leaves open the possibility that the redistribution of the membrane proteins may involve actin-dependent endocytosis and intracellular trafficking using cytoplasmic microtubule motors.

Although visualizing NAP1 has proven difficult, the actin ring structures that appear after prolonged Lat B treatment are likely comprised of NAP1 filaments, as *nap1* mutant cells are unable to form phalloidin-labeled rings upon Lat B treatment (**Figure 7G**). Given the transient nature of the actin rings, it is possible that these rings acutely compensates for actin-dependent functions to protect the nucleus. These functions may include regulating nuclear shape (Khatau et al., 2009), nuclear positioning (Gundersen and Worman, 2013), or connecting with the nucleoskeleton to transduce mechanical signals (Tapley and Starr, 2013). Alternatively, the ring may aid in transport to or from the nuclear pore complexes (Mosalaganti et al., 2018), where we see clusters of filaments gather prior to Lat B disruption (**Figure 3C**).

A seemingly universal function of actin is directing mitotic and meiotic events by localizing vesicles and the cytokinetic machinery in plant, fungi, and animal cells. Filamentous actin-rich ring structures have been associated with cytokinesis in animal cells (Wang, 2005), yeast (Arai and Mabuchi 2002), and recently, it was reported that an actin shell ruptures the nucleus to mediate meiosis in starfish oocytes (Wesolowska et al., 2018). Many of these rings undergo a constriction that is mediated by a contractile class II myosin. Given that *Chlamydomonas* cells lack this type of myosin and use an understudied microtubule-dependent structure called the phycoplast for cytokinesis (Johnson and Porter, 1968; Holmes and Dutcher, 1989; Gaffel and el-Gammel, 1990; Schibler and Huang, 1991), actin may play a more minor role in cell division in this organism relative to others.

The co-expression of a conventional and divergent actin within a single cell establishes *Chlamydomonas* as an excellent new system to identify fundamental principles of actin biology. Robust visualization of the filamentous actin network in these cells may reveal novel modes of actin-dependent regulation for major cellular processes, such as ciliary biology and photosynthesis, for which *Chlamydomonas* is already a leading model organism.

## Acknowledgements

We would like to thank Masayuki Onishi, John Pringle, and Fred Cross for providing the *nap1* mutant. Thank you to members of the Avasthi Lab for troubleshooting and manuscript feedback. We would like to acknowledge Wolfgang Baumeister and Jurgen Plitzko for enabling the cryo-ET work by providing support and instrumentation. This work was funded through P20GM104936 and P20GM103418 (to P.A.), the Deutsche Forschungsgemeinschaft grant EN 1194/1-1 as part research unit FOR2092 (to B.D.E.), and The Max Planck Society.

